# Insights on cross-species transmission of SARS-CoV-2 from structural modeling

**DOI:** 10.1101/2020.06.05.136861

**Authors:** João PGLM Rodrigues, Susana Barrera-Vilarmau, João MC Teixeira, Elizabeth Seckel, Panagiotis Kastritis, Michael Levitt

**Author notes:** These authors contributed equally to this work (Susana Barrera-Vilarmau, João MC Teixeira).

## Abstract

Severe acute respiratory syndrome coronavirus 2 (SARS-CoV-2) is responsible for the ongoing global pandemic that has infected more than 14 million people in more than 180 countries worldwide. Like other coronaviruses, SARS-CoV-2 is thought to have been transmitted to humans from wild animals. Given the scale and widespread geographical distribution of the current pandemic, the question emerges whether human-to-animal transmission is possible and if so, which animal species are most at risk. Here, we investigated the structural properties of several ACE2 orthologs bound to the SARS-CoV-2 spike protein. We found that species known not to be susceptible to SARS-CoV-2 infection have non-conservative mutations in several ACE2 amino acid residues that disrupt key polar and charged contacts with the viral spike protein. Our models also predict affinity-enhancing mutations that could be used to design ACE2 variants for therapeutic purposes. Finally, our study provides a blueprint for modeling viral-host protein interactions and highlights several important considerations when designing these computational studies and analyzing their results.

## Introduction

SARS-CoV-2, a novel betacoronavirus first identified in China in late 2019, is responsible for the ongoing global pandemic that has infected more than 14 million people worldwide and killed over 600.000 [1]. Comparative genomics studies suggest that SARS-CoV-2 was transmitted to humans from an animal host, most likely bats or pangolins [2]. Given the widespread human-to-human transmission across the globe, the question emerges whether humans can infect other animal species with SARS-CoV-2, namely domestic and farm animals. Identifying potential intermediate hosts that can act as reservoirs for the virus has both important global health, animal welfare, and ecological implications.

During the course of this pandemic, there have been several news reports of domestic, farm, and zoo animals testing positive for SARS-CoV-2 infection. Belgium [3] and New York [4] reported positive symptomatic cases in cats, The Netherlands reported infection of minks in farms [5], and the Bronx Zoo in New York reported infections in lions and tigers [6]. In all these cases, the vehicle of transmission appears to be an infected human owner or handler. More importantly, in the case of the mink farms in The Netherlands, there is evidence of human-to-animal-to-human transmission. In addition to these reported cases, several groups put forward both pre-prints and peer-reviewed studies on animal susceptibility to SARS-CoV-2 under controlled laboratory conditions [7–9], two of which are of particular interest. The first study showed that cats, civets, and ferrets are susceptible to infection; pigs, chickens, and ducks are not, while the results for dogs were inconclusive [7]. A second study, using human cells expressing recombinant SARS-CoV-2 receptor proteins showed that camels, cattle, cats, horses, sheep, and rabbit can be infected with the virus, but not chicken, ducks, guinea pigs, pigs, mice, and rats [8]. Together, these studies provide a dataset of confirmed susceptible and non-susceptible species that we can analyze to find molecular discriminants between the two groups. For simplicity, from here on we will refer to susceptible and non-susceptible species as SARS-CoV-2^pos^ and SARS-CoV-2^neg^, respectively.

Like SARS-CoV-1 before, SARS-CoV-2 infection starts with the binding of the viral spike protein to the extracellular protease domain of angiotensin-converting enzyme 2 (ACE2) [10], a single-pass transmembrane protein expressed on the surface of a variety of tissues, including along the respiratory tract and the intestine. Several biophysical and structural studies identified helices α1 and α2, as well as a short loop between strands β3 and β4 in ACE2 as the interface for the viral spike protein [10–13]. These studies also identified key differences between the sequences of the receptor binding domains (RBD) of SARS-CoV-1 and SARS-CoV-2, which explain the stronger interaction of the latter with human ACE2. If binding to ACE2 is the first step in the infection cycle, we can reasonably assume that sequence variation across ACE2 orthologs can explain why only some animal species are susceptible to infection. In addition, combining structural and binding data with the natural diversity of ACE2 across species can help elucidate the key aspects that drive ACE2 interaction to viral RBDs and ultimately help guide the development of therapeutic molecules against SARS-CoV-2.

Unsurprisingly, several groups already contributed multiple sequence and structure-based analyses of how sequence variation affects ACE2 binding to SARS-CoV-2 RBD [14–17]. Two recent preprints, specifically, focus on the effects of ACE2 variation on RBD binding. The first used an ACE2 sequence library to select for mutants that bind RBD with high affinity, identifying several mutants that enhance or decrease affinity to the viral protein and providing a blueprint for engineering proteins and peptides with therapeutic purposes [14]. While useful, we note that the authors carried out a single round of selection as opposed to the multiple rounds commonly carried out in similar studies. The second study used computational modeling to predict ΔΔG of mutations in 215 animal species and assess their risk for infection [15]. In addition, the authors also identified a number of locations on ACE2 that contribute to binding the viral RBD, in particular residues 31, 38, 353, as well as a cluster of N-terminal hydrophobic amino acid residues.

In this study, we aimed to leverage structural, binding, and sequence data to investigate how different ACE2 orthologs bind to SARS-CoV-2 RBD. We selected 29 animal species likely to encounter humans in a variety of residential, industrial, and commercial settings. For each of these species, we generated 3D models of ACE2 bound to RBD and refined these models using short molecular dynamic simulations. After refinement, we found that models of SARS-CoV-2^pos^ species generally have a lower (better) score than those of SARS-CoV-2^neg^ species. Further, we carried out a per-residue energy analysis that predicts both key locations in ACE2 that are consistently mutated across SARS-CoV-2^neg^ species, as well as possible mutations that likely enhance binding to the viral RBD. Collectively, our results provide a structural framework to understand why certain animal species are not susceptible to SARS-CoV-2 infection, and also provide a starting point for rational engineering of antiviral molecular therapeutics. Finally, our work also provides a blueprint for future studies of viral-host protein interactions at high-resolution.

## Results

All our models and the scoring statistics are available for visualization and download at https://github.com/joaorodrigues//ace2-animal-models/.

### Sequence conservation of ACE2 orthologs

We analyzed the sequence conservation of ACE2 across our dataset, with respect to the entire sequence (591 residues) and to the interface residues computed from a structure of ACE2 bound to RBD (PDB ID: 6m17) (22 residues) (Table 1). All orthologs are reasonably conserved, with global similarity values to the human ACE2 sequence (hACE2) ranging from 72% (goldfish) to 99.5% (chimpanzee) (S1 Fig). All species coarsely cluster in three classes consistent with evolutionary distance to humans: primates have the highest similarity values, followed by other mammals, birds and reptiles, and finally fish. Zooming in on the interface residues, we find more variation (Fig 1, left). Similarity values for this region range from 50% (crocodile) to 100% (all 3 primates) but, despite an overall correlation (Pearson R^2^ of 0.69), they do not always match global similarities. Hedgehogs and sheep, for example, share 86.7% and 86.4% global similarity with hACE2, respectively, but 59% and 95.5% for the interface region. In other words, sheep share 21 out of 22 residues with hACE2 at the interface with RBD, while hedgehogs share 13. The horseshoe bat, one of the proposed animal reservoirs for SARS-CoV-2, shares 72.2% interface similarity with hACE2, a comparable value to the 77.3% of the SARS-CoV-2^neg^ mouse sequence. Altogether, these results prompt two observations. First, neither global nor interface sequence similarity is predictive of SARS-CoV-2 susceptibility. Second, that the interface of the viral RBD is substantially plastic and able to bind to sufficiently different ACE2 orthologs.

**Fig 1.**
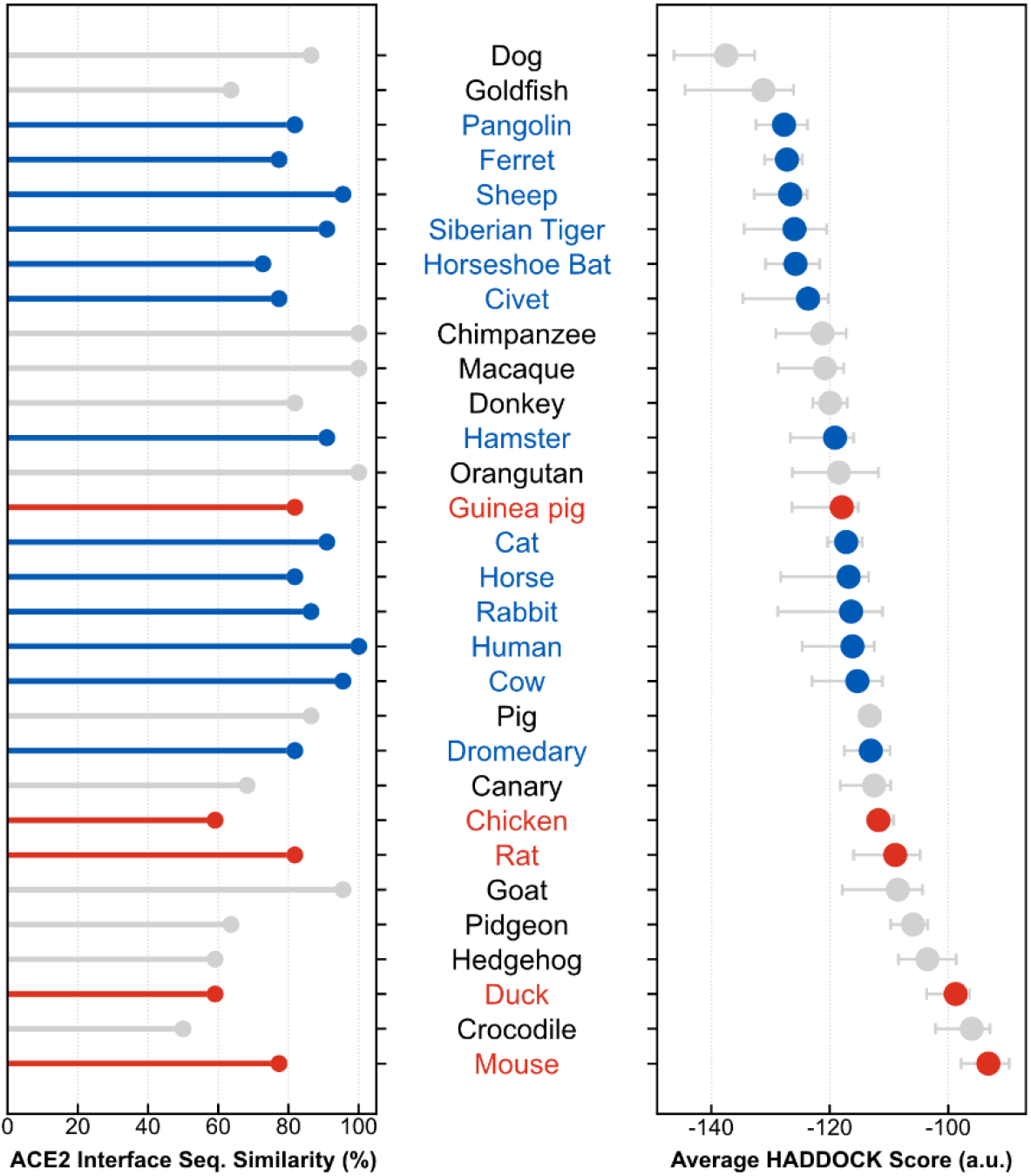
Interface statistics of modeled ACE2:RBD complexes. SARS-CoV-2^pos^ species (in blue) generally have lower (better) HADDOCK scores (left, expressed in arbitrary units) than SARS-CoV-2^neg^ species (in red). A similar but less conclusive trend is observed between the sequence similarities of amino acid residues interacting with the viral RBD (derived from PDB 6m17) (right). Collectively, these results suggest that SARS-CoV-2^neg^ species lack specific key ACE2 amino acid residues, leading to impaired binding between the two proteins. Species are ordered in increasing order of HADDOCK score. Species for which SARS-CoV-2 susceptibility is unknown or assays were inconclusive are shown in gray.

**Table 1.** List of species included in the study

| Scientific Name | Common Name | NCBI Protein ID |
| --- | --- | --- |
| *Homo sapiens* | Human | NP_001358344.1 |
| *Anas platyrhynchos* | Duck | XP_012949915.2 |
| *Bos taurus* | Cow | XP_005228485.1 |
| *Camelus dromedarius* | Dromedary | XP_010991717.1 |
| *Canis lupus familiaris* | Dog | NP_001158732.1 |
| *Capra hircus* | Goat | NP_001277036.1 |
| *Carassius auratus* | Goldfish | XP_026131313.1 |
| *Cavia porcellus* | Guinea pig | XP_023417808.1 |
| *Columba livia* | Pidgeon | XP_021154486.1 |
| *Crocodylus porosus* | Crocodile | XP_019384826.1 |
| *Equus asinus* | Donkey | XP_014713133.1 |
| *Equus caballus* | Horse | XP_001490241.1 |
| *Erinaceus europaeus* | Hedgehog | XP_007538670.1 |
| *Felis catus* | Cat | XP_023104564.1 |
| *Gallus gallus* | Chicken | XP_416822.2 |
| *Macaca mulatta* | Macaque | NP_001129168.1 |
| *Manis javanica* | Pangolin | XP_017505746.1 |
| *Mesocricetus auratus* | Hamster | XP_005074266.1 |
| *Mus musculus* | Mouse | NP_081562.2 |
| *Mustela putorius furo* | Ferret | NP_001297119.1 |
| *Oryctolagus cuniculus* | Rabbit | XP_002719891.1 |
| *Ovis aries* | Sheep | XP_011961657.1 |
| *Paguma larvata* | Civet | AAX63775.1 |
| *Pan troglodytes* | Chimpanzee | XP_016798468.1 |
| *Panthera tigris altaica* | Siberian Tiger | XP_007090142.1 |
| *Pongo abelii* | Orangutan | NP_001124604.1 |
| *Rattus norvegicus* | Rat | NP_001012006.1 |
| *Rhinolophus sinicus* | Horseshoe bat | AGZ48803.1 |
| *Serinus canaria* | Canary | XP_009087922.1 |
| *Sus scrofa* | Pig | NP_001116542.1 |

### Refinement of the hACE2:RBD complex

In order to validate the refinement protocol used in our analysis, we created and refined models of human ACE2 (hACE2) bound to SARS-CoV-2 RBD. We used the cryo-EM structure of full-length human ACE2 bound to the RBD, in the presence of the amino acid transporter B^0^AT1 (PDB ID: 6m17). Compared to a high-resolution crystal structure of the same complex (PDB ID: 6m0j), the cryo-EM structure lacks several key contacts between our two proteins of interest, which we attribute to poor density for side-chain atoms at the interface region. Our refinement protocol restores the majority of these contacts (S1 Table), yielding an average HADDOCK score of −116.2 (arbitrary units, a.u.) for the 10 best models of the best cluster. See Materials and Methods for further details on the protocol. These scores fall within the range observed for a reference set of transient protein-protein interactions (N=144, HADDOCK score=-124.9 ± 53.4) [18]. Upon visual inspection, the interfaces in our models are dominated by hydrogen bond interactions involving the ACE2 α1 helix and a small loop between strands β3 and β4. There is one single salt-bridge involving hACE2 D30 and RBD K417 consistently present in all our hACE2 models. These observations all agree with the published crystal structure. Further, the buried surface area of the refined models is also in agreement with published crystal structures (~1800 Å^2^). As such, we are confident that our modeling and refinement protocol is robust enough to model all ACE2 orthologs.

### Refinement of orthologous ACE2:RBD complexes

We modeled and refined complexes for all 29 ACE2 orthologs in our dataset (Table 1) using the same protocol as above. The representative models for each species (10 best models of the best cluster) are available for visualization and download at https://joaorodrigues.github.io/ace2-animal-models/. The HADDOCK scores of all 30 ACE2 complexes (including hACE2) range from −137.5 (dog) to −93.2 (mouse), indicating substantial differences between these interfaces (Fig 1, right, and S2 Table). The average HADDOCK score is −116.4, very close to that of the human complex (−116.2). Overall, models of SARS-CoV-2^pos^ species have consistently lower (better) scores than those of SARS-CoV-2^neg^ species. Although it is well-known that docking scores do not quantitatively correlate with experimental binding affinities [19], these scores suggest that SARS-CoV-2^neg^ species lack one or more key ACE2 residues that contribute significantly to the interaction with RBD.

To understand what forces drive the interactions between ACE2 and SARS-CoV-2 RBD, we quantified the contribution of each component of the HADDOCK scoring function to the overall score (Fig 2). The HADDOCK score is a linear combination of van der Waals, electrostatics, and desolvation energy terms. In our models, electrostatics are the most discriminatory component (Pearson R^2^ of 0.60), followed by desolvation (0.31), and finally van der Waals (0.08). These correlations suggest that differences between the models of the different species originate primarily in polar and charged residues, in agreement with observations from experimental structures. In addition, the buried surface area of the models also correlates quite strongly with the HADDOCK score (Pearson R^2^ of 0.66), which is unsurprising since larger interfaces tend to make more contacts. Most models bury between 1700 and 1850 Å^2^, in agreement with the crystal and cryo-EM structures, while the top-scoring species (dog and goldfish) bury nearly 2000 Å^2^ and the lowest-scoring (mouse) bury only 1600 Å^2^. Finally, there is a weak correlation between the average HADDOCK score of the representative models and the sequence similarity of the ACE2 interface residues (Pearson R^2^ of 0.18) (S2 Fig).

**Fig 2.**
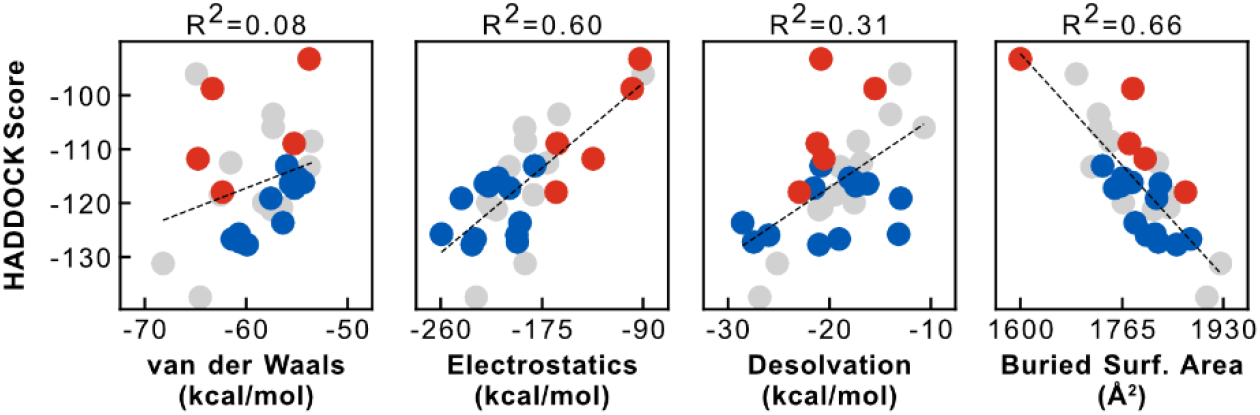
Correlation of HADDOCK score with individual energy terms and structural features. Differences in electrostatics energy contribute the most towards discriminating SARS-CoV-2^pos^ species (blue) from SARS-CoV-2^neg^ species (red), supporting observations of hydrogen bonding networks and charged interactions in experimental structures. The buried surface area of the models is also strongly correlated with their HADDOCK score, suggesting larger interfaces of SARS-CoV-2^pos^ species might confer better binding properties.

### Structural and energetic differences between SARS-CoV-2^pos^ and SARS-CoV-2^neg^ species

To gain further insight on how ACE2 sequence variation across the different orthologs affects binding to SARS-CoV-2 RBD, we calculated HADDOCK scores for each interface residue in the refined models. This high-resolution analysis highlights several ACE2 amino acids with strong interaction energies that differ between SARS-CoV-2^pos^ and SARS-CoV-2^neg^ species (Figs. 3 and S3).

**Fig 3.**
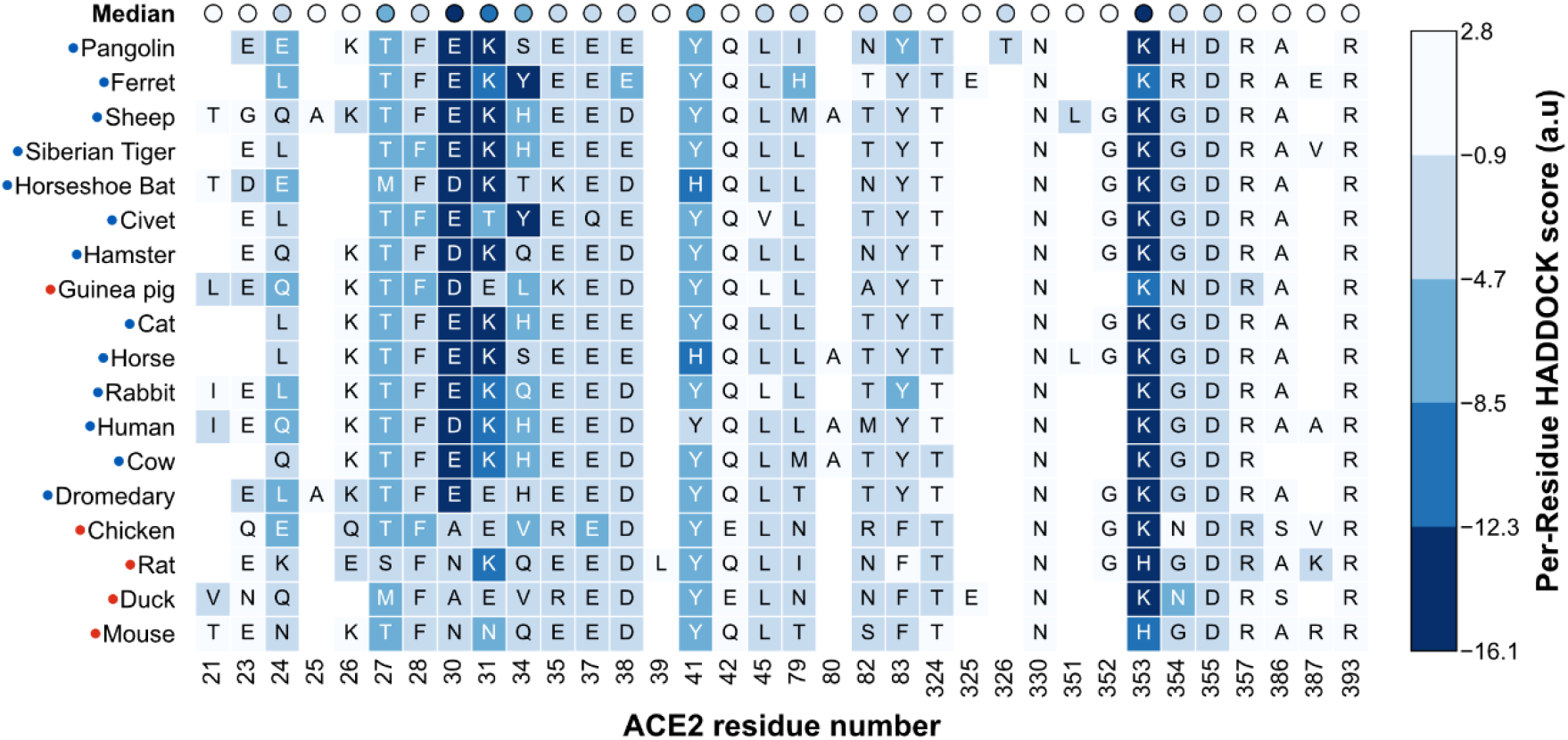
HADDOCK score of individual ACE2 interface residues. For each species (row), the blocks (columns) represent amino acid residues within 5Å of the viral RBD in any of the species’ best 10 models. The identity of the amino acid is shown in one-letter code. The colors represent the HADDOCK score of each residue, averaged over the 10 models: lower scores (dark blue) indicate more favorable interactions, while positive scores indicate steric clashes or electrostatic repulsion. The first row shows the median of the averages for each column. From this analysis, we predict that amino acid residues at positions 30, 31, and 353 contribute the most to the stability of the ACE2:RBD complex. In SARS-CoV-2^neg^ species (red labels), some of these residues are consistently mutated (30 and 31), which could explain their lower susceptibility to infection. S3 Fig shows the per-residue analysis for all species in the dataset.

The first and most relevant of these sites is amino acid 30, which in hACE2 (D30) interacts with RBD K417 to form the only intermolecular salt-bridge of the interface (Fig 4, top left). In all 12 SARS-CoV-2^pos^ species, this site is occupied by a negatively charged amino acid residue. In contrast, 4 out of 5 SARS-CoV-2^neg^ species have a hydrophobic or polar residue at this position, breaking the intermolecular salt-bridge (Fig 4, bottom left). The second site is amino acid 31, a lysine in hACE2, and in nearly all of the SARS-CoV-2^pos^ species, that interacts both with ACE2 E35 and RBD Q493 (Fig 4, top middle). The only exceptions are the civet and dromedary sequences, mutated to threonine and glutamate, respectively. In the case of the civet, our models show that T31 can still hydrogen bond with both E35 and RBD Q493. Dromedaries, on the other hand, share E31 with chickens, guinea pigs, and ducks, all SARS-CoV-2^neg^ species. However, and quite beautifully, in dromedary ACE2 the likely electrostatic repulsion between E31 and E35 is compensated by a lysine at position 76 (Q76 in hACE2) leading to the formation of an additional intramolecular salt-bridge that possibly stabilizes the fold of ACE2 and frees E35 to hydrogen bond with Q493 (90% of our models). Those three SARS-CoV-2^neg^ species have an additional charge-reversal mutation at position 35. In all our chicken and duck models, E31 is locked in an intramolecular salt-bridge with R35, weakening the intermolecular hydrogen bond with RBD Q493 (Fig 4, bottom middle). Finally, guinea pigs compensate K31E with E35K and remain able to hydrogen bond with RBD, while rats have a lysine at this position.

**Fig 4.**
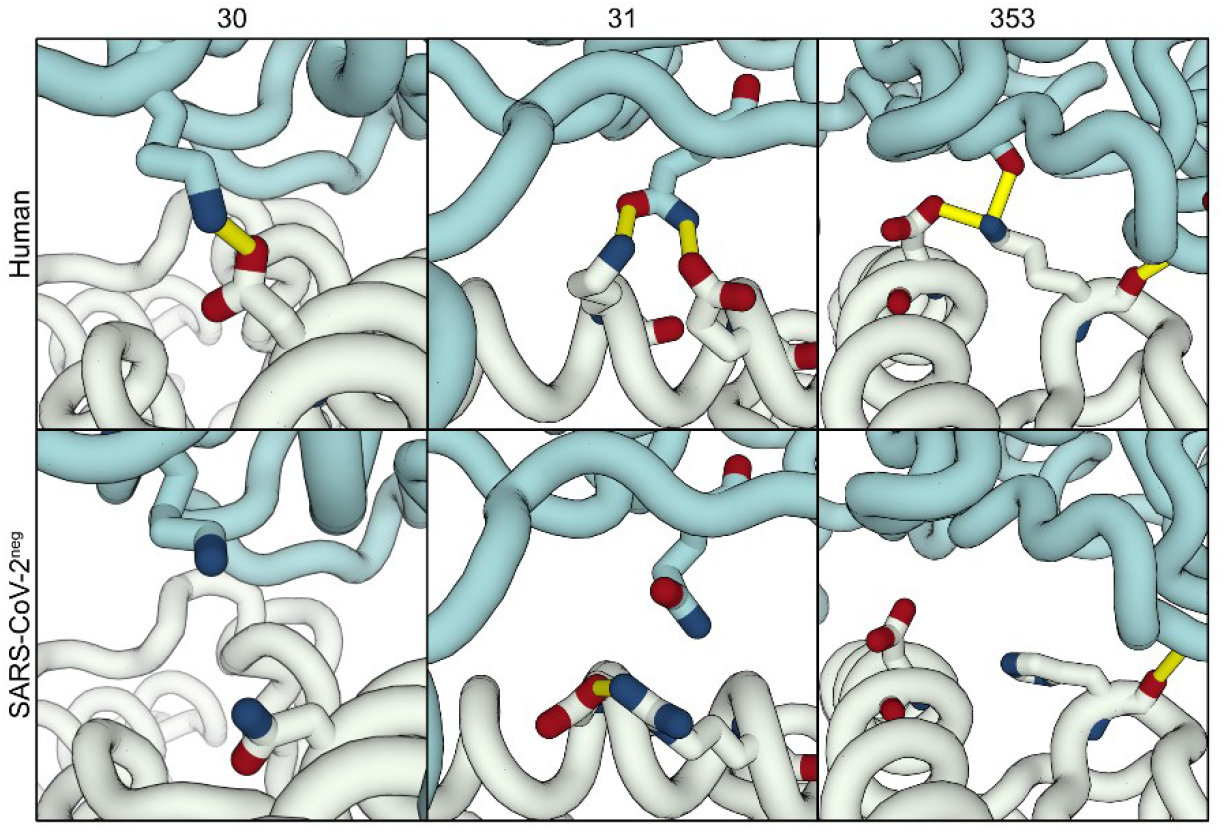
Interface differences between human and SARS-CoV-2^neg^ models. The top panels show key residue-residue interactions at the interface between hACE2 (white) and the viral RBD (teal), which are conserved in nearly all SARS-CoV-2^pos^ species: salt-bridge between D30 and K417 (left); three-body interaction between K31, E35, and RBD Q493 (middle); and the interactions of K353, an intramolecular salt-bridge with D38 and an intermolecular hydrogen bonds with G496 and G502 (right). The bottom panels highlight equivalent regions in three SARS-CoV-2^neg^ species: D30N mutation in mice (left) disrupts the intermolecular salt-bridge; D31K/D35R in ducks stabilizes an intramolecular salt-bridge and weakens the intermolecular hydrogen bond (middle); K353H in mice disrupts the intramolecular salt-bridge (right).

Besides these major discriminatory sites, we identified multiple other sites that are systematically mutated in SARS-CoV-2^neg^ species. The first of these sites is K353 (in hACE2), which is involved in an intramolecular salt-bridge with D38, and two hydrogen bonds with RBD G496 and G502 (Fig 4, bottom right). In rat and mouse ACE2, both SARS-CoV-2^neg^ species, this residue is mutated to a histidine, which weakens the interaction with D38, possibly leading to increased conformational dynamics of the β3-β4 loop and consequently lower binding affinity. Then, Q42, conserved in most other species, hydrogen bonds with RBD Y449 in the majority of our models. In canary, chicken, pigeon, hedgehog, duck, and crocodile ACE2 sequences, this amino acid is mutated to a glutamate, which introduces the possibility of an additional intramolecular salt-bridge with K68, in ACE2 helix α2. As we observe in some of our models, this intramolecular interaction prevents the formation of the intermolecular hydrogen bond. Finally, amino acid 83, a tyrosine in hACE2 and all other SARS-CoV-2^pos^ is mutated to phenylalanine in 4 out of 5 SARS-CoV-2^neg^ species: mouse, duck, rat, and chicken. The loss of the hydroxyl group excludes residue 83 from a ternary hydrogen-bonding network involving Q24 and RBD N487 that likely stabilizes the protein-protein interface. Without this hydrophilic terminal group, residue 83 might also prefer less solvent accessible conformations in the unbound state, burying between both α1 and α2 helices and thus being less available to interact with RBD F486.

### Affinity-enhancing mutations from ACE2 orthologs

In addition to highlighting discriminatory mutations between SARS-CoV-2^pos^ and SARS-CoV-2^neg^ species, our models also allow us to search for mutations that could be used to generate variants of hACE2 with higher affinity towards the viral RBD. To this end, we calculated a modified HADDOCK score for each residue, including both intra- and intermolecular interactions, and then subtracted the score of the corresponding residue in the hACE2:RBD models (see Material and Methods for details) (Fig 5).

**Fig 5.**
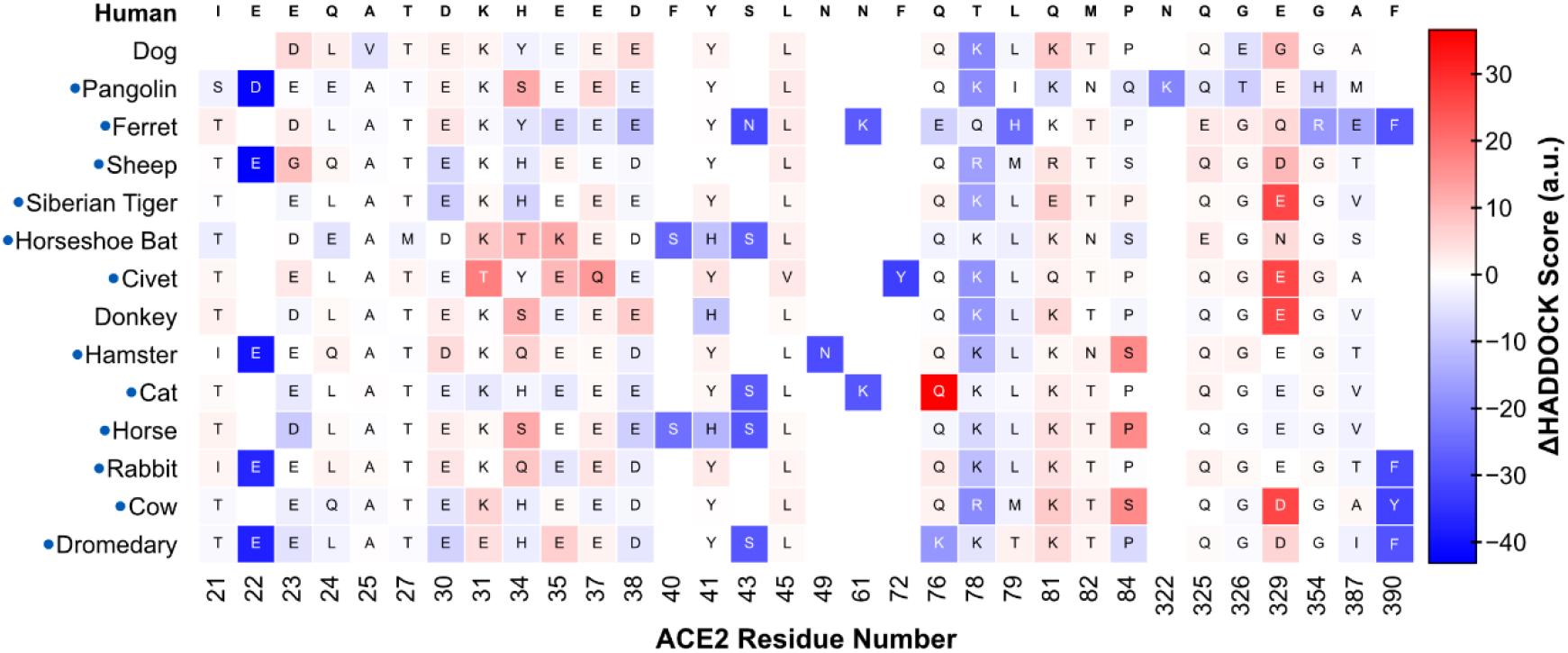
ΔHADDOCK score of individual ACE2 interface residues compared to hACE2. For each species (row), the blocks (columns) represent amino acid residues within 7.5Å of the viral RBD in any of the species’ best 10 models. The identity of the amino acid is shown in one-letter code. The colors represent the ΔHADDOCK score – including intramolecular interactions – of each residue, averaged over the 10 models, compared to the average of the corresponding hACE2 residue: negative scores (dark blue) indicate a stabilizing mutation. This analysis highlights several potential affinity-enhancing mutations, namely Q24E, A25V, D30E, H34Y, F40S, Y41H, F72Y, L79H, and A387E. We note that this analysis requires further visual inspection of the models to account for additional variations in ACE2 sequence that may skew the per-residue HADDOCK score. Refer to the main text for details. S4 Fig shows the same plot for all species of the dataset.

The resulting analysis highlights several single-point mutations that we predict could confer a higher affinity for RBD if engineered on hACE2. Some we can explain with simple biophysics following a careful inspection of the models (Fig 6). Q24E, observed in both the pangolin and horseshoe bat sequences, contributes to a stronger hydrogen bond network with partner RBD N487, and helps stabilize the α1 helix through interactions with the backbone of neighboring S21; A25V, observed only in the dog sequence, is buried between helices α1, α2, and α3, and contributes to a stronger packing with neighboring hydrophobic and aromatic residues (L29, Y83, V93, and L97); D30E stabilizes the intermolecular salt-bridge with RBD K417 due to the longer glutamate side-chain; H34Y enhances the hydrophobic interactions with neighboring L455 and the aliphatic chain of RBD N493; F72Y introduces possible hydrogen-bonding interactions between helices α1 and α2, while maintaining strong hydrophobic packing through the phenyl ring; L79H, observed only in the ferret ACE2 sequence, allows for intermolecular hydrogen bonds with the backbone carbonyl of RBD G485, in addition to stabilizing helix α2 and the packing of helices α1 and α2 through hydrogen bonds with residue 76 and aromatic stacking with F28; finally, A387E, observed in the ferret sequence, can interact with both R354 (G354 in hACE2) and, more importantly, RBD R408.

**Fig 6.**
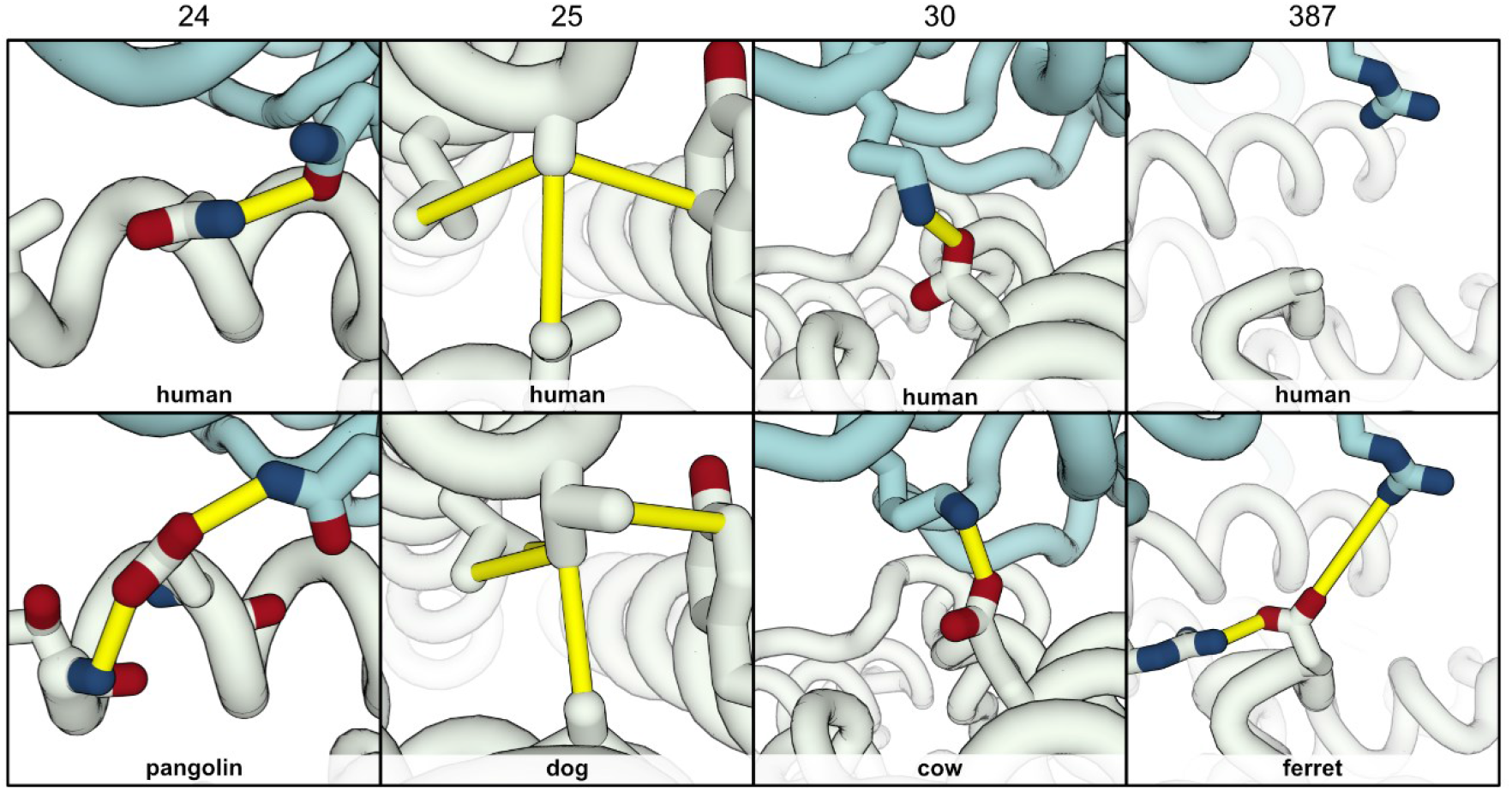
Predicted affinity-enhancing mutations for hACE2. Analyzing the residue energetics of ACE2 orthologs suggests mutations that have the potential to enhance the affinity of hACE2 (white) to RBD (teal). The top panels shows our top-scoring hACE2:RBD model and its interactions (yellow cylinders) for four such sites: residues 24, 25, 30, and 387. The bottom panels show mutations in specific species, and the resulting new or enhanced interactions: Q24E in pangolin, A25V in dog, D30E in cow, and A387E in ferret. Some of these mutations are found in multiple ACE2 orthologs.

Other mutations observed in top-scoring species and predicted in our models to have stronger local interactions are dependent on additional mutations in neighboring residues. F40S, observed in bat and horse ACE2, forms a hydrogen bond with the hydroxyl group of Y390 (F390 in hACE2); Y41H, observed in bat, donkey, and horse ACE2, contributes to a polar network involving RBD residues, namely Q498, T500, and N501, as well as hydrogen bonds with the backbone of ACE2 L351, that might help stabilize the local fold of the β3-β4 loop; lastly, Q76E, Q76K, and T78K all stabilize helices α1 and α2 through interactions with adjacent residues, such as E31 (dromedary), E74 (pangolin), E75 (dog), or H79 (ferret).

## Discussion

### Can structural modeling predict cross-species transmission of SARS-CoV-2?

Our computational modeling of 30 vertebrate ACE2 orthologs bound to SARS-CoV-2 RBD discriminates between previously reported SARS-CoV-2^pos^ and SARS-CoV-2^neg^ species. Models of SARS-CoV-2^neg^ species – chicken, duck, mouse, and rat – have clearly higher (worse) HADDOCK scores than average (Fig 1), suggesting that these species’ non-susceptibility to infection could stem from deficient RBD binding to ACE2. Despite this clear trend, our predictions are not entirely correct. Our modeling ranks guinea pig ACE2 (SARS-CoV-2^neg^) as a better receptor for SARS-CoV-2 RBD than for example, human, cat, horse, or rabbit ACE2 (all SARS-CoV-2^pos^ species), despite experiments showing that there is negligible binding between the two proteins [8].

These results highlight the need for carefully evaluating predictions from computational models. As noted earlier in the introduction, SARS-CoV-2 infection is a complex multi-step process [20]. Thus, while we can assume that impaired ACE2 binding decreases odds of infection, we cannot state that ACE2 binding is predictive of infection. For instance, experiments with recombinant ACE2 show that the pig ortholog binds SARS-CoV-2 RBD and leads to entry of the virus in host cells [8], but tests in live animals returned negative results [7]. Our modeling protocol, like any structure prediction method, is limited by the accuracy of its sampling method (how many conformational states do we model) and of its scoring function (how do we pick the most native-like model). In addition, by basing our models on a cryo-EM structure of the hACE2:RBD complex, we make assumptions about the bound state of the two proteins when it is known from structures of the full-length SARS-CoV-2 spike protein [21] and coarse-grained simulations [22] that there is considerable flexibility at the interface. As such, our computational models alone cannot be used to predict whether certain animal species are at risk of infection; our ranking of species needs thorough experimental validation. What our models do predict, however, are distinctive molecular features between SARS-CoV-2^pos^ and SARS-CoV-2^neg^ species. As the adage goes, ‘all models are wrong, but some are useful.’

### SARS-CoV-2^neg^ species lack important polar and charged ACE2 residues

Having established that our modeling protocol and scoring function generally distinguishes between SARS-CoV-2^pos^ and SARS-CoV-2^neg^ species, we took a closer look at the biophysical properties of the interface across all orthologs. We decomposed our scoring function in its individual energy terms (Fig 2) and found that SARS-CoV-2^neg^ models rank worse due to a substantial decrease in electrostatic energy and interface surface area. An exhaustive per-residue energy analysis of the 10 best models for each species (300 models in total), reveals a loss of hydrogen bonds and salt-bridges in models of mouse, duck, rat, and chicken ACE2. Most importantly, these species lack the only intermolecular salt-bridge observed between hACE2 and RBD due to mutations on position 30 (D30 in hACE2) (Fig 4, left). These predictions are supported by experimental work, where mutants lacking a negative charge at this position are largely unable to bind SARS-CoV-2 RBD [14]. Non-conservative mutations at other sites on ACE2 also contribute negatively to the interface scores. Our models suggest that the introduction of a negatively charged residue at position 31 is disruptive to binding, in agreement with experiments [14], unless it is compensated by an additional mutation. In both chicken and duck ACE2, the compensatory mutation – E35R – nevertheless locks E31 in an intramolecular salt-bridge and prevents it from interacting with RBD (Fig 4, middle). Then, at the opposite end of the α1 helix, our models identify K353 as a strong contributor to interface stability that is mutated in both rat and mouse ACE2 (Fig 4, right). The long lysine side-chain stabilizes the interface region of ACE2 by forming an intramolecular salt-bridge with D38, but also contributes to hydrogen bonds with viral RBD, with G496 and G502. These results support other modeling work [17] that predicts that RBD mutants G496D bind worse to ACE2 because of the disruption of this intramolecular salt-bridge. Also, as shown by experiments, any mutation in this region decreases the ability of ACE2 to bind RBD [14], confirming our predictions and highlighting a second important interaction site lacked by rodents’ ACE2. Our predictions identify other sequence differences between SARS-CoV-2^pos^ and SARS-CoV-2^neg^ species that impair intra- and intermolecular polar interactions. Position 83 is mutated from a tyrosine to a phenylalanine in all SARS-CoV-2^neg^ species except guinea pig, while position 42, a glutamine in all SARS-CoV-2^pos^ species, is mutated to a glutamate in chicken and duck ACE2. Introducing these mutations on hACE2 leads to impaired binding of RBD [14]. Given the highly polar nature of the interface (Tables S1 and S2), it is then plausible that the accumulation of several mutations on key polar and charged residues, as observed in SARS-CoV-2^neg^ species, leads to a drastic reduction in binding affinity between the two proteins.

### Natural variants of ACE2 encode potential affinity-enhancing mutations for SARS-CoV-2 RBD

One of the many proposed antiviral agents against SARS-CoV-2 is recombinant soluble hACE2 [23], which acts as a decoy for the viral RBD and therefore reduces rates of infection. While deep mutagenesis experiments have been used to optimize protein-protein interfaces for therapeutic purposes [24], it is impractical to carry out an exhaustive search of the entire protein sequence space. Our models suggest several sites and variants that potentially enhance the affinity between hACE2 and RBD (Fig 5). These predictions fall in three broad categories: the stabilization of existing interactions, the introduction of novel interactions, and stabilization of the ACE2 interface region. We note, however, that our coverage of sequence space is limited to naturally occurring variants, and that natural selection imposes additional constraints on sequence variability besides RBD binding.

For the first category, the clearest affinity enhancer seems to be D30E, a variant observed in 6 of the 8 best scoring species (Fig 6, third panel) and shown in experiments to increase binding to RBD [8,14]. The longer side-chain of a glutamate residue can help strengthen and stabilize the intermolecular salt-bridge with RBD K417. The impact of such Asp-to-Glu mutations in modulating protein interactions has been reported previously for other systems [25]. Other mutations predicted to enhance the strength of existing interactions between ACE2 and RBD include Q24E (Fig 6, first panel) and F72Y, both validated by experiments [14]. The introduction of novel interactions is particularly interesting from a protein design perspective. Our models predict that placing a negative charge at position 387 might allow for a second intermolecular salt-bridge to form with RBD R408 (Fig 6, fourth panel). In our hACE2 models, RBD R408 points towards – but does not interact with – the glycan molecule bound to N90. It has been shown that removing this N-glycosylation motif increases RBD binding, while both A387D and A387E lead to mild increases in binding affinity in some cases [14]. As such, we propose that a double N90A/A387E mutant could have a synergistic effect on RBD affinity. Finally, it is known that interactions between rigid binders, with little to none conformational changes upon binding, have the highest affinities [18]. Indeed, this is a ground rule of many successful protein interface designs (e.g. [26]). Our per-residue energy analysis predicts that A25V stabilizes the packing of the α1 and α2 helices, which is an important nexus of RBD interactions (Fig 6, second panel).

Our models also predict that mutating H34Y increases RBD binding, possibly by introducing novel interactions with RBD via the terminal hydroxyl of the tyrosine side-chain. In addition, the large aromatic ring offers a hydrophobic partner for RBD L455. Our predictions for both H34V and H34S indicate that neither of these mutants is energetically favorable, likely because they retain only one of the two types of interactions (aromatic or polar). However, experiments show exactly the opposite behavior: H34S or H34V dramatically increase binding to RBD, while H34Y decreases it [14]. This result highlights the limitations of our models and stresses the need for experimental validation for all our predictions.

In summary, our protocol combines structural, sequence, and binding data to create a structure-based framework to understand SARS-CoV-2 susceptibility across different animal species. Our models help rationalize the impact of naturally-occurring ACE2 mutations on SARS-CoV-2 RBD binding and explain why certain species are not susceptible to infection with the virus. In addition, we propose possible affinity-enhancing mutants that can help guide engineering efforts for the development of ACE2-based antiviral therapeutics. Our protocol and models can easily be replicated using freely-available tools and web servers and serve as a blueprint for future modeling studies on protein interactions where data is available for a large number of homologues.

Finally, to prevent human-to-animal transmission, we recommend following the World Organization for Animal Health guidelines: people infected with COVID-19 should limit contact with their pets, as well as with other animals (including humans).

## Materials and Methods

### Sequence Alignment of ACE2 Orthologs

Sequences of ACE2 orthologs from 28 species were retrieved from NCBI using the human gene as a reference (Gene ID: 59272, updated on 20-Apr-2020) and the query term “ortholog_gene_59272[group]”. Other species, such as *Rhinolophus sinicus*, were manually included using custom queries. The sequences were aligned with MAFFT version 7 [27,28], using the alignment method FFT-NS-i (Standard). Some sequences had undefined amino acids (‘X’), which we converted to glycine to allow modeling without any bias for amino acid identity. All species and the respective protein identifiers are listed in Table 1.

### Definition of Sequence Similarity

All calculations were based on the alignments from MAFFT, restricted to the region used for modeling (residues 21-600). To calculate sequence similarity, we considered the following groups based on physico-chemical properties: charged-positive (Arg, Lys, His), charged-negative (Asp, Glu), aromatic (Phe, Tyr, Trp), polar (Ser, Thr, Asn, Gln), and apolar (Ala, Val, Ile, Met). Cys, Gly, and Pro residues were considered individual classes.

### Modeling of ACE2 Orthologs

The modeling of ACE2 orthologs was carried out using MODELLER 9.24 [29] and custom Python scripts (available here: https://github.com/joaorodrigues//ace2-animal-models/). We used the cryo-EM structure of the SARS-CoV-2 RBD bound to human ACE2 (PDB ID: 6M17) [12] as a template for all our subsequent models, including all glycans and the coordinates of RBD. To save computational resources, we modeled only the extracellular domain of ACE2, specifically residues 21-600, which are known to be sufficient to bind to RBD. To avoid unwanted deviation from the initial cryo-EM structure, we restricted the optimization and refinement of the models to the coordinates of atoms of mutated or inserted residues. We used the *fastest* library schedule for model optimization and the *very_fast* schedule for model refinement. For each species, we generated 10 backbone or loop models and selected the one with the lowest normalized DOPE score as a representative. These final models were then processed to remove any sugar molecules in species where the respective asparagine residue had been mutated.

### Refinement of ACE2:RBD complexes

The initial complex models were prepared for refinement using the pdb-tools suite [30]. Each chain was separated into a different PDB file (pdb_selchain) and standardized with TER and END statements (pdb_tidy). We used HADDOCK 2.4 [31] to carry out the refinement of the models. The protein molecules were parameterized using the standard force field in HADDOCK, while the sugars were parameterized using updated parameters for carbohydrates [32]. We used a modified version of the topology generation scripts to allow automatic detection of N-linked glycans and expand the range of the interface refinement (10 Å distance cutoff). Each initial homology model was refined through 50 independent short molecular dynamics simulations in explicit solvent (solvshell=True). These refined models were then clustered using the FCC algorithm [33] with default parameters and scored using the HADDOCK score, a linear combination of van der Waals, electrostatics, and desolvation. A lower HADDOCK score is better. The top 10 models of the top scoring cluster, ranked by its average HADDOCK score, were selected as representatives of the complex.

### Analysis of interface contacts of refined ACE2:RBD complexes

We used the interfacea analysis library (version 0.1) (http://doi.org/10.5281/zenodo.3516439) to identify intermolecular contacts between hACE2 and RBD, specifically hydrogen bonds, salt bridges, and aromatic ring stacking. Hydrogen bonds were defined between any donor atom (nitrogen, oxygen, or sulfur bound to a hydrogen atom) within 2.5 Å of an acceptor atom (nitrogen, oxygen, or sulfur), if the donor-hydrogen-acceptor angle was between 120 and 180 degrees. Salt bridges were defined between two residues with a pair of cationic/anionic groups within 4 Å of each other. Finally, two aromatic residues were defined as stacking if the centers of mass of the aromatic groups were within 7.5 Å (pi-stacking) or 5 Å (t-stacking) and the angle between the planes of the rings was between 0 and 30 degrees (pi-stacking) or between 60 and 90 degrees (t-stacking). Additionally, for pi-stacking interactions, the projected centers of both rings must fall inside the other ring. For each modelled species, we took the 10 best models of the best cluster, judged by their HADDOCK score, and aggregated all their contacts together. Contacts present in at least 5 models were considered representative.

### Per-residue decomposition of HADDOCK scores

We used a custom CNS [34] script to calculate the HADDOCK score of each residue at the interface between ACE2 and RBD. Briefly, the protocol was the following. For each model, since HADDOCK uses a united-atom force field, we first added missing hydrogen atoms and minimized their coordinates, keeping all other atoms fixed. We marked a residue of ACE2 as part of the interface if any of its atoms were within 5 Å of any atom of RBD, and *vice-versa*. We then calculated the electrostatics, van der Waals, and desolvation energies for each of these residues, considering only atoms belonging to the other protein chain. Note that this protocol does not account for intramolecular effects of mutations. Finally, we calculated the HADDOCK score per residue, using the default scoring function weights, and averaged per-residue values for the best 10 models of the best cluster of each species. For the calculation of combined intra- and intermolecular scores (Fig 6), we followed a similar protocol where the distance cutoff to define neighbors was increased to 7.5 Å and atoms from both chains were considered.

## Acknowledgements

The authors thank R. Almeida, T. Dots, M. Farzan, K. Lindorff-Larsen, J. Puglisi, and R. Fernandes for feedback and encouragement during the course of the project.

## Supporting Information

**Fig. S1.**
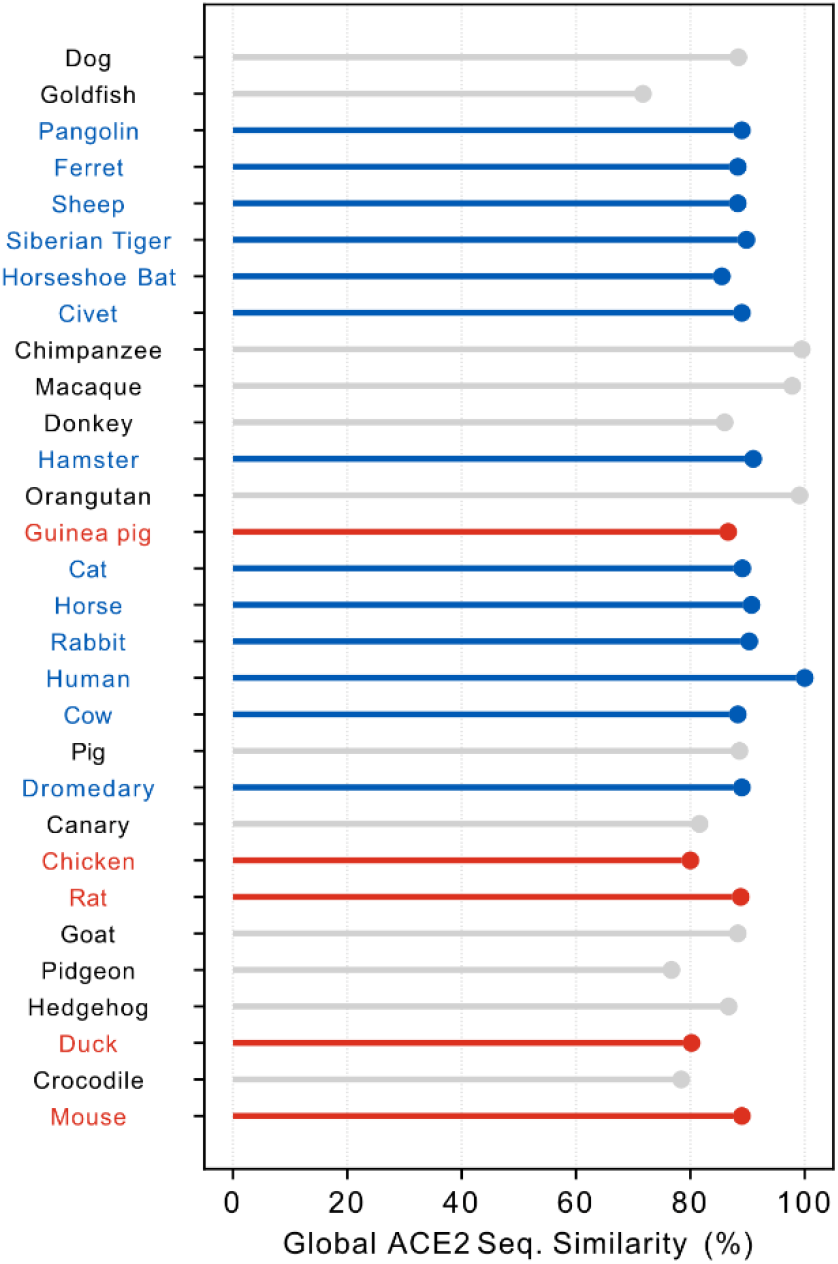
Global sequence similarity across ACE2 orthologs.

**Fig. S2.**
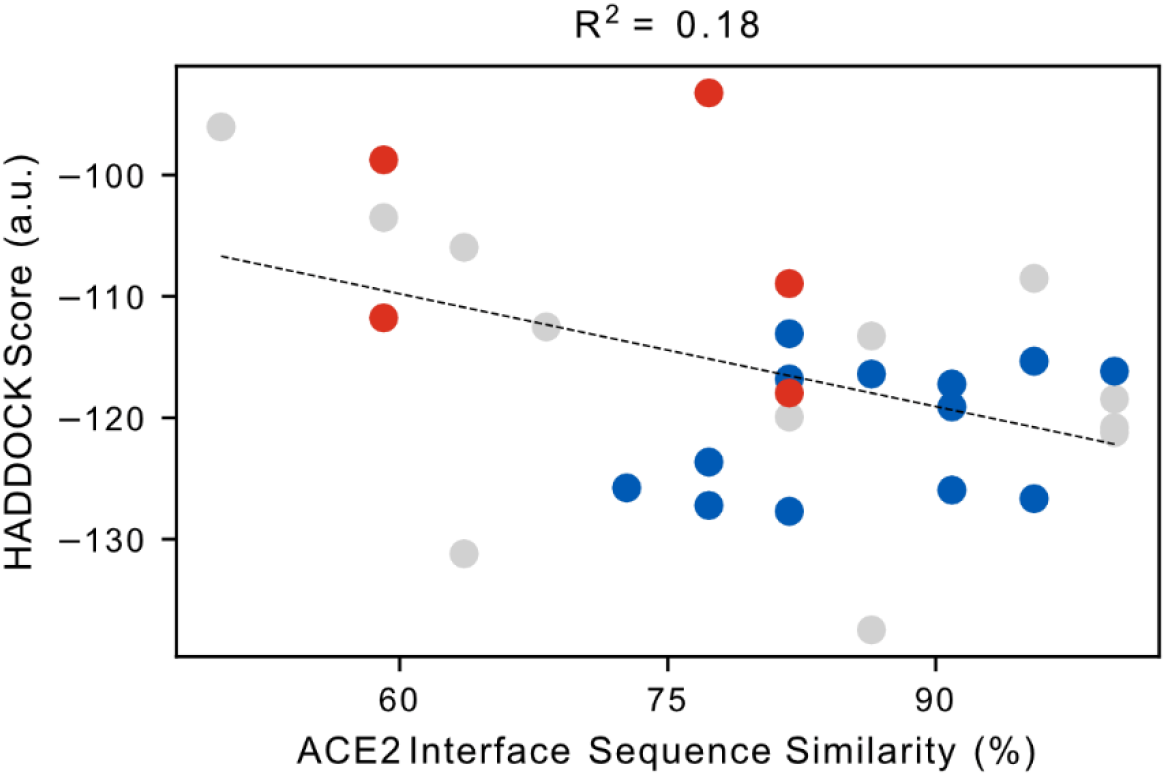
Correlation between HADDOCK score and interface sequence similarity for all models.

**Fig. S3.**
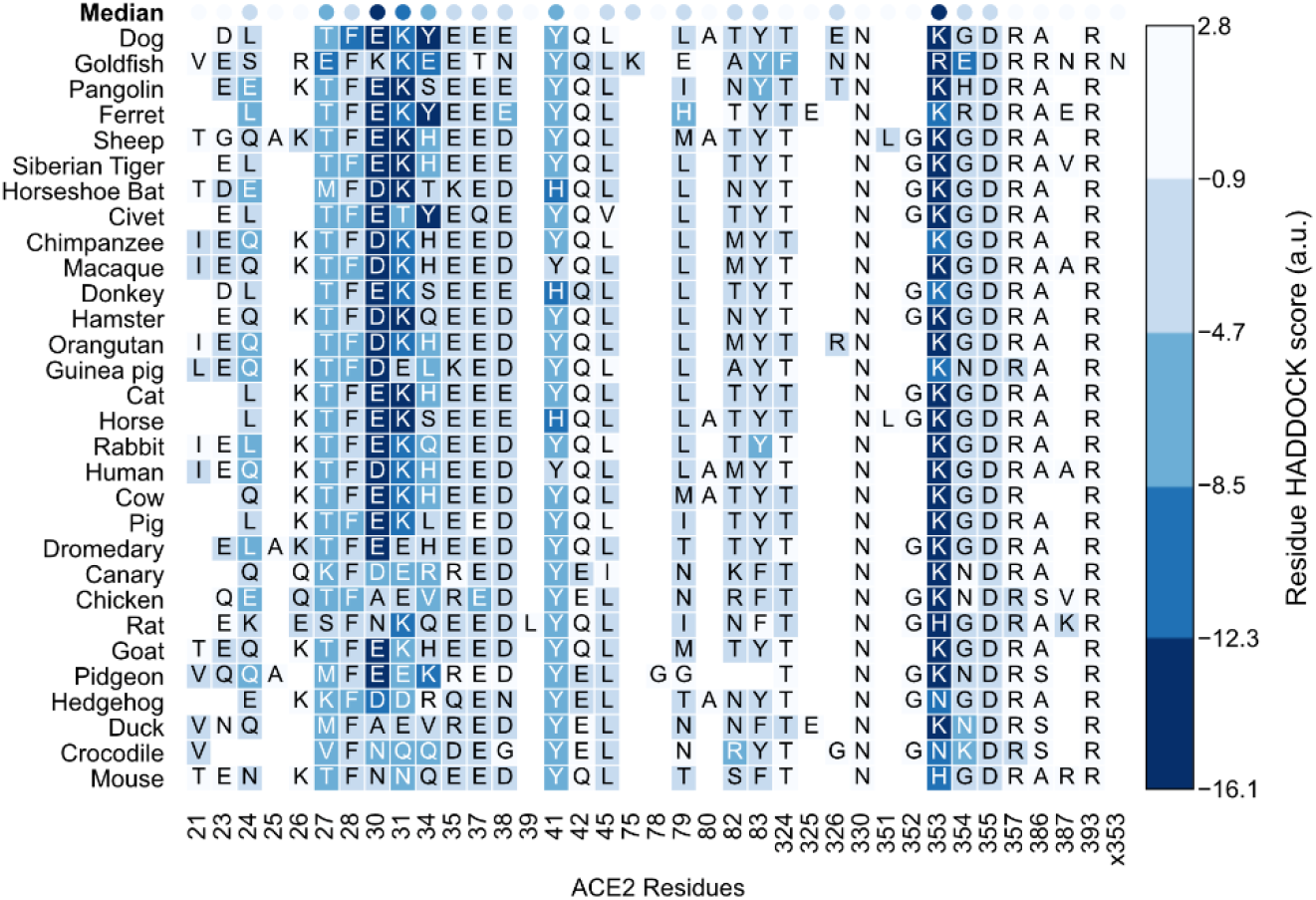
HADDOCK score of individual ACE2 interface residues for all species.

**Fig. S4.**
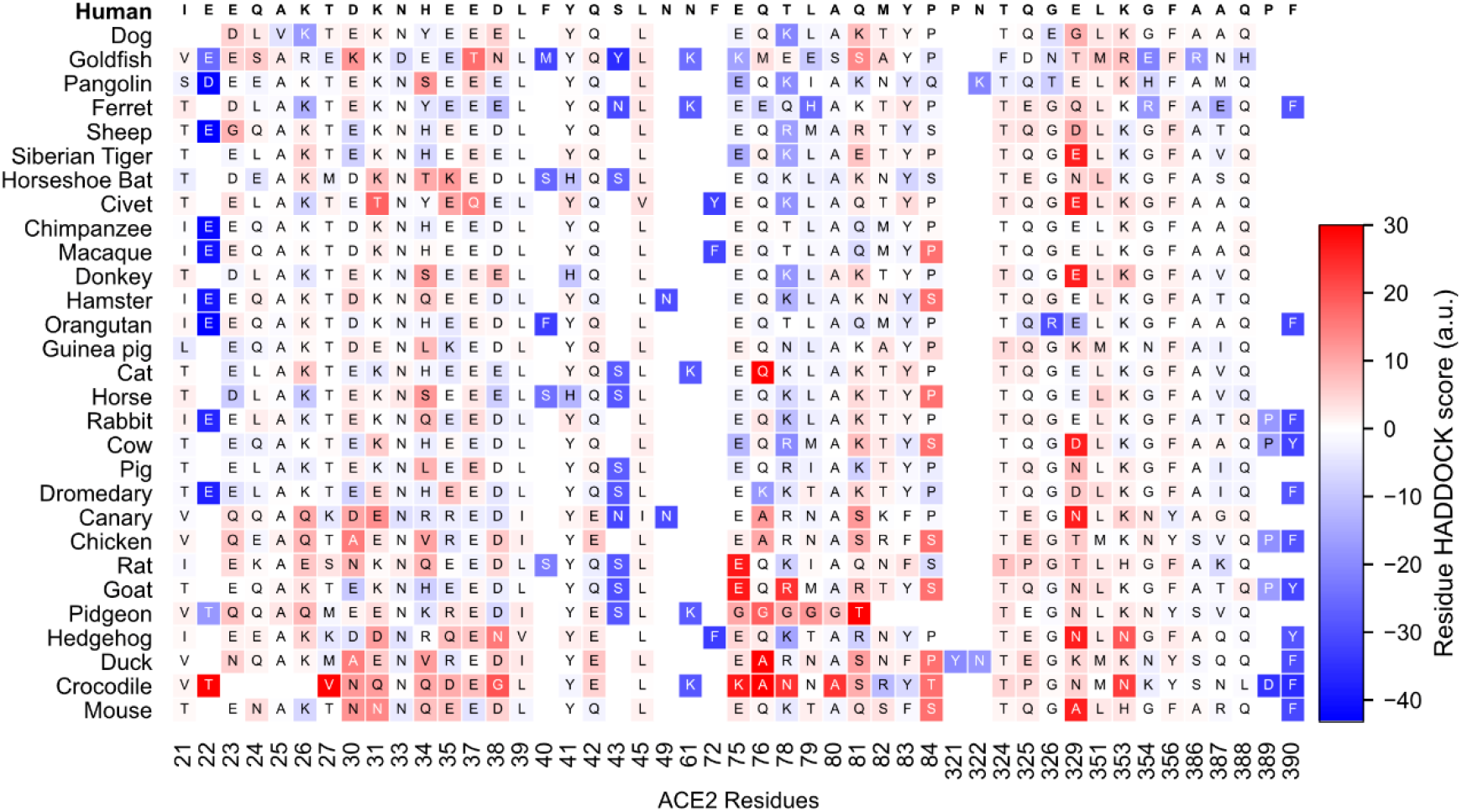
ΔHADDOCK score of individual ACE2 interface residues compared to hACE2 for all species.

**Table S1.** Interface contacts of refined human ACE2:RBD

| Type | ACE2 Residue | RBD Residue | Frequency in Top 10 hACE2 Models | In crystal (6m0j) |
| --- | --- | --- | --- | --- |
| hbond | Q24 | A475 | 7 |  |
| hbond | Q24 | N487 | 6 | x |
| hbond | T27 | Y489 | 9 |  |
| ionic | D30 | K417 | 10 | x |
| hbond | K31 | F490 | 4 |  |
| hbond | K31 | Q493 | 10 | x |
| hbond | H34 | Y453 | 2 |  |
| hbond | E35 | Q493 | 6 | x |
| hbond | E37 | Y505 | 4 | x |
| hbond | D38 | Y449 | 10 | x |
| hbond | Y41 | T500 | 10 | x |
| hbond | Y83 | N487 | 2 | x |
| hbond | K353 | Y495 | 1 |  |
| hbond | K353 | G496 | 9 |  |
| hbond | K353 | G502 | 10 | x |

**Table S2.** HADDOCK scores and individual energy terms for each modeled ACE2:RBD complex. The values represent the average and standard deviation of the 10 best models (ranked by HADDOCK score) of each species.

| Species | HADDOCK Score (a.u.) | van der Waals (kcal/mol) | Electrostatics (kcal/mol) | Desolvation (a.u.) | Buried Surface Area (Å²) |
| --- | --- | --- | --- | --- | --- |
| Dog | -137,5 ± 3,7 | -64,5 ± 4,0 | -230,3 ± 18,4 | -26,9 ± 2,5 | 1903 ± 44 |
| Ferret | -127,2 ± 1,8 | -60,6 ± 2,6 | -195,8 ± 16,8 | -27,5 ± 3,4 | 1824 ± 70 |
| Goldfish | -131,2 ± 5,9 | -68,2 ± 3,8 | -189,3 ± 19,2 | -25,1 ± 3,2 | 1925 ± 67 |
| Pangolin | -127,7 ± 3,3 | -59,9 ± 3,6 | -233,8 ± 16,0 | -21,0 ± 2,0 | 1854 ± 32 |
| Hamster | -119,1 ± 3,1 | -57,6 ± 2,8 | -242,8 ± 23,8 | -13,0 ± 2,5 | 1821 ± 51 |
| Siberian Tiger | -126,0 ± 4,8 | -60,7 ± 3,9 | -196,3 ± 27,8 | -26,0 ± 3,3 | 1804 ± 37 |
| Guinea pig | -118,0 ± 3,3 | -62,3 ± 2,6 | -163,3 ± 12,2 | -23,0 ± 2,5 | 1868 ± 41 |
| Sheep | -126,7 ± 2,5 | -61,4 ± 2,2 | -231,2 ± 26,8 | -19,0 ± 3,6 | 1876 ± 25 |
| Chimpanzee | -121,2 ± 3,4 | -57,5 ± 3,7 | -213,5 ± 17,6 | -21,0 ± 3,1 | 1816 ± 33 |
| Civet | -123,6 ± 4,2 | -56,4 ± 4,5 | -193,6 ± 27,9 | -28,6 ± 3,4 | 1786 ± 56 |
| **Human** | -116,2 ± 3,2 | -54,4 ± 2,8 | -221,8 ± 15,1 | -17,5 ± 3,2 | 1781 ± 37 |
| Dromedary | -113,1 ± 2,2 | -56,0 ± 2,0 | -181,3 ± 16,3 | -20,8 ± 2,7 | 1733 ± 65 |
| Horseshoe bat | -125,8 ± 2,7 | -60,7 ± 3,2 | -259,4 ± 17,5 | -13,2 ± 3,2 | 1819 ± 52 |
| Pig | -113,3 ± 1,1 | -53,8 ± 2,1 | -202,7 ± 12,6 | -18,9 ± 2,5 | 1715 ± 32 |
| Cow | -115,3 ± 4,1 | -54,9 ± 1,5 | -212,1 ± 19,1 | -18,0 ± 4,0 | 1766 ± 41 |
| Cat | -117,2 ± 1,7 | -55,2 ± 2,0 | -202,4 ± 9,8 | -21,5 ± 3,0 | 1752 ± 42 |
| Macaque | -120,8 ± 3,2 | -56,5 ± 2,1 | -217,7 ± 18,6 | -20,7 ± 2,3 | 1841 ± 37 |
| Pidgeon | -106,0 ± 2,0 | -57,3 ± 2,7 | -189,7 ± 11,9 | -10,7 ± 3,0 | 1730 ± 28 |
| Orangutan | -118,5 ± 4,5 | -62,5 ± 2,7 | -182,3 ± 21,1 | -19,5 ± 3,8 | 1837 ± 49 |
| Horse | -116,8 ± 4,2 | -55,4 ± 3,7 | -220,0 ± 14,0 | -17,4 ± 2,4 | 1762 ± 34 |
| Rabbit | -116,4 ± 5,6 | -55,6 ± 2,7 | -222,6 ± 18,2 | -16,2 ± 2,7 | 1827 ± 55 |
| Donkey | -119,9 ± 1,9 | -58,3 ± 1,7 | -220,5 ± 13,5 | -17,6 ± 3,3 | 1767 ± 44 |
| Goat | -108,5 ± 4,0 | -53,5 ± 3,6 | -189,0 ± 10,5 | -17,2 ± 3,1 | 1745 ± 64 |
| Chicken | -111,8 ± 1,4 | -64,8 ± 1,6 | -132,3 ± 15,0 | -20,5 ± 3,7 | 1802 ± 44 |
| Canary | -112,5 ± 2,8 | -61,5 ± 5,0 | -170,7 ± 33,7 | -16,8 ± 3,8 | 1822 ± 73 |
| Hedgehog | -103,5 ± 3,2 | -57,3 ± 5,3 | -160,9 ± 38,9 | -14,0 ± 3,8 | 1726 ± 48 |
| Rat | -108,9 ± 3,5 | -55,3 ± 4,5 | -162,4 ± 16,0 | -21,2 ± 2,7 | 1777 ± 58 |
| Crocodile | -96,0 ± 2,5 | -64,9 ± 2,7 | -90,4 ± 18,1 | -13,1 ± 2,6 | 1690 ± 49 |
| Duck | -98,8 ± 2,3 | -63,3 ± 1,9 | -99,5 ± 6,0 | -15,5 ± 3,1 | 1782 ± 55 |
| Mouse | -93,2 ± 2,6 | -53,8 ± 2,6 | -93,1 ± 14,4 | -20,8 ± 2,5 | 1598 ± 65 |

## Author Contributions

**Conceptualization:** João P. G. L. M. Rodrigues

**Funding Acquisition:** Michael Levitt

**Investigation:** João P. G. L. M. Rodrigues, Susana Barrera-Vilarmau, João M. C. Teixeira, Panagiotis Kastritis

**Writing – Original Draft:** João P. G. L. M. Rodrigues, João M. C. Teixeira, Susana Barrera-Vilarmau, Elizabeth Seckel

**Writing – Review and Editing:** João P. G. L. M. Rodrigues, Susana Barrera-Vilarmau, João M. C. Teixeira, Elizabeth Seckel, Panagiotis Kastritis

